# Variation and diversification of the microbiome of *Schlechtendalia chinensis* on two alternate host plants

**DOI:** 10.1101/351882

**Authors:** Haixia Wu, Xiaoming Chen, Hang Chen, Qin Lu, Zixiang Yang, Weibin Ren, Juan Liu, Shuxia Shao, Chao Wang, Kirst King-Jones, Ming-Shun Chen

## Abstract

*Schlechtendalia chinensis*, a gall-inducing aphid, has two host plants in its life cycle. Its wintering host is a moss (typically *Plagiomnium maximoviczii*) and its main host is *Rhus chinensis* (Sumac), on which it forms galls during the summer․. This study investigated bacteria associated with *S. chinensis* living on the two different host plants by sequencing 16S rRNAs. A total of 183 Operational Taxonomic Units (OTUs) from 50 genera were identified from aphids living on moss, whereas 182 OTUs from 49 genera were found from aphids living in Sumac galls. The most abundant bacterial genus among identified OTUs from aphids feeding on both hosts was *Buchnera*. Despite similar numbers of OTUs, the composition of bacterial taxa showed significant differences between aphids living on moss and those living on *R. chinensis*. Specifically, there were 12 OTUs from 5 genera (family) unique to aphids living on moss, and 11 OTUs from 4 genera (family) unique to aphids feeding in galls on *R. Chinensis*. Principal Coordinate Analysis (PCoA) also revealed that bacteria from moss-residing aphids clustered differently from aphids collected from galls. Our results provid a foundation for future analyses on the roles of symbiotic bacteria in plant - aphid interactions in general, and how gall-specific symbionts differ in this respect.

## Introduction

Insects harbour a wide range of symbiotic microbes, but the actual species composition can vary between different developmental stages [1, 2]. In general, bacterial symbionts are comprised of two kinds: obligate symbionts and facultative symbionts. Obligate endosymbionts are necessary for insect growth and development, and usually supply important nutrients such as essential amino acids to the host insect [3, 4]. In contrast, facultative symbionts are not essential for growth and development of their host, but may improve fitness. For instance, some facultative symbionts boost the defense system of the host and thus enhance resistance to natural enemies and pathogens [5–7]. Typically, facultative symbionts have not shared a long evolutionary relationship with their host insects [8].

Aphids are a large group of insects belonging to the Aphidiodea superfamily. Many aphid species are destructive pests of crops and forests. Symbiosis with microorganisms is critical for the development of aphids. One of the essential symbiotic bacteria associated with aphids is *Buchnera aphidicola*, an obligate endosymbiont that provides essential amino acids, which are either absent or scarce in plant sap [9]. In addition to its obligate symbionts, aphids can also harbor several facultative bacterial symbionts, which can be mutualistic in the context of various ecological interactions. *Serratia symbiotica* is one of the most common facultative symbiotic bacteria in aphids [10]. S. *symbiotica*, together with other symbionts, provides aphids with protection against parasitoids [11]. Because of different ecological habitats, different aphid species harbor specific bacterial taxa for specific functions [12].

The aphid *Schlechtendalia chinensis*, a member of the subfamily Pemphiginae, is a beneficial species with great economic and medicinal values [13, 14]. The aphid induces the formation of large, single-chambered galls on its host plant, and some chemicals of the galls are used as ingredientsin traditional Chinese medicine to treat various conditionsincluding cough, diarrhea, night sweats, dysentery, and internal bleeding [15]. In the mining industry, some applications use chemicals from *S. chinensis*-induced galls to extract rare metals. In addition to its economic and medicinal values, *S. chinensis* also has a fascinating biology. The aphid has a life cycle of sexual and asexual reproduction, and a switch of host plant species during its life cycle. The primary host of *S. chinensis* is *Rhus chinensis* (Sumac tree) and the secondary host is *Plagiomnium maximoviczii* (moss). In early spring in China, sexuparae migrate from mosses to the trunk of Sumac trees to produce sexually reproductive females and males that subsequently mate to produce fundatrix, which feed on rachide wings and induce the formation of galls [16, 17]. Within a gall, *S. chinensis* undergoes cyclical parthenogenesis for multiple generations to increase population sizes rapidly. In late fall, the galls break up and the alate fundatrigeniae of *S. chinensis* emerge fromgalls and relocateto their secondary host mosses, where aphids over-winter.

Based on its unique biology and its ability to induce galls, we hypothesized that *S. chinensis* harbors a set ofunique symbiontic taxa, thatare likely to play critical roles in its biology, such as mediating adaptive responses on different hosts, or contributing toits gall-inducing ability. The rapid advance in high throughput sequencing technologies has made it possible to comprehensively investigate the compositions of microbial community in small insects [18]. In this study, we examined the diversity of bacterial communities associated with *S. chinensis* in different hosts via next generation sequencing. Specifically, we sequenced the variable regions 4-5 of 16S rRNA amplicons using the Illumina sequencing platform (using barcoded Illumina paired-end sequencing, BIPES) to characterize the composition and diversity of microbiomes during different developmental stages of *S. chinensis.*

## Materials and methods

### Aphid collection

We collected aphid samples corresponding to nine time points (one per month) directly from host plants from March to November, 2015, in Kunming, Yunnan province, China. Each condition was represented by three independent replicates. In order to observe potential dynamic changes in the diversity of microbal taxa, samples were collected every month, and for consistency, we chose the middle of each month. The April samples represented the fundratrix stage․. Samples from March, April, and November were taken as the group of insects from the host moss, whereas the remaining samples were taken from Sumac trees.

To obtain insects from galls, individual galls were soaked in 75% ethanol by gently shaking for 30 s. The gall was washed three times with doubly distilled (dd) water. Using sterile conditions, an incision into the gall was carried outwith a blade. Aphids were taken out individually with forceps and transferred into individual tubes. After removing any environmental contaminants, the insect samples were frozen at −80 °C for subsequent genomic DNA extraction. For moss-residing aphids, 30 individual aphids were collected and transferred into a microfuge tube. Aphids were washed with 1 ml of 75% ethanol by gently shaking for 60 s. The insects were then stored in −80 °C for DNA extraction.

### Extraction of genome DNA

Total genomic DNA from a sample was extracted using a QIAampDNA Micro Kit (OMEGA Forensic DNA Kit) following the directionsprovided by the manufacturer. DNA concentration and purity was monitored on a 1% agarose gel. DNA concentration was diluted to 1ng/μL with sterile water and used as template for PCR amplification.

### Amplificon generation

The variable regions 4-5 (16SV4-V5) in the internal transcribed spacers (ITS) of 16S rRNA genes were amplified using a pair of specific primers, namely 515F “5- GTGCCAGCMGCCGCGG-3” and 907R”5- CCGTCAATTCMTTTRAGTTT-3”,with barcodes. All PCR reactions were carried out with Phusion^®^ High-Fidelity PCR Master Mix (New England Biolabs). A PCR reaction mix contained 5 μ;l 5× reaction buffer, 5 μ;l 5× GC buffer, 2 μl dNTP (2.5 mM), 1 μl Forward primer and Reverse primer, respectively (10 mM), 2 μl DNA Template, 8.75 μl ddH2O, and 0.25 μl Q5 DNA Polymerase. PCR amplification was achieved by an initial denaturation at 98°Cfor 2min, followed by 25-30 cycles of denaturation (98° for 15s), annealing (55° for 30s), and extension (72° for 30s). A final extension step at 72° for 5 min was included at the end of each PCR amplification.

### PCR amplification and sequencing

PCR products were mixed with the same volume of 1x loading buffer (containing SYBR green) and analyzed on a 2% agarose gel. Samples with a main 400-450bp band were purified with a Qiagen Gel Extraction Kit (Qiagen, Germany). Amplicons were then quantified and sequencing libraries were generated using NEB Next Ultra DNA library Prep Kit for Illumina (New England Biolabs Inc. USA) according to manufacturer’s recommendations. The amplicon libraries were subsequently sequenced on an Illumina MiSeq platform at Personalbio (Shanghai, P.R. China).

### Bioinformatics analyses

Paired-end reads were merged using FLASH [19]. Low quality reads were removed. Sequences were analyzed with the QIIME software package [20] using default parameters for each step. The Uparse software [21] was used to cluster the sequences into operational taxonomic units (OTUs) at an identity threshold of 97%. The RDP Classifier [22] was used to assign each OTU to a taxonomic level. Other analyses, including rarefaction curves, Shannon index, and Good’s coverage, were performed with QIIME. In addition, the OTU table produced by the QIIME pipeline was imported into MEGAN 4 and mapped on the GreenGene Database [23, 24].

Principal Coordinate Analysis (PCoA) was performed to get principal coordinates and visualize from complex, multidimensional data. PCoA has been recognized as a simple and straight-forward method to group and separate samples in a dataset. In this study, PCoA was used to analyze the sequencing results using WGCNA package, stat packages and ggplot2 package in R software.

### Statistical methods

All experimental results were evaluated using analysis of variance (ANOVA) for multiple comparisons followed by the Turkey test. Differences were considered significant at p < 0.05. All analyses were done using the programs WGCNA, STAT, and ggplot2 in the R software package.

## Results

### Sequence reads and OTUs

In total, 975,583 raw reads were obtained from all samples, and after filtering, 973,373 high-quality sequences were retained and subsequently clustered into 194 OTUs (S1 Table). Of the 194 OTUs, five were classified as Archaea, 83 were Bacteria, and the remaining 106 remain to be determind since they showed no blast hits. Of the 82 bacterial OTUs, 56 (67.5%) were Proteobacteria, 10 belong to Firmicutes, and the remaining 17 belong to Cyanobacteria and others.

The distribution of reads and OTUs duringdifferent seasons is shown in Table 1, along with other parameters including Chao1 Index, Shannon Index, and Good’s coverage. The highest number of reads was obtained in the sample collected on October 19, but the highest number of OTUs was obtained in the sample collected on March 24. In comparison, the lowest number of reads was obtained in the sample collected on May 27, whereas the lowest number of OTUs was obtained in the sample collected on September 16.

**Table 1.**
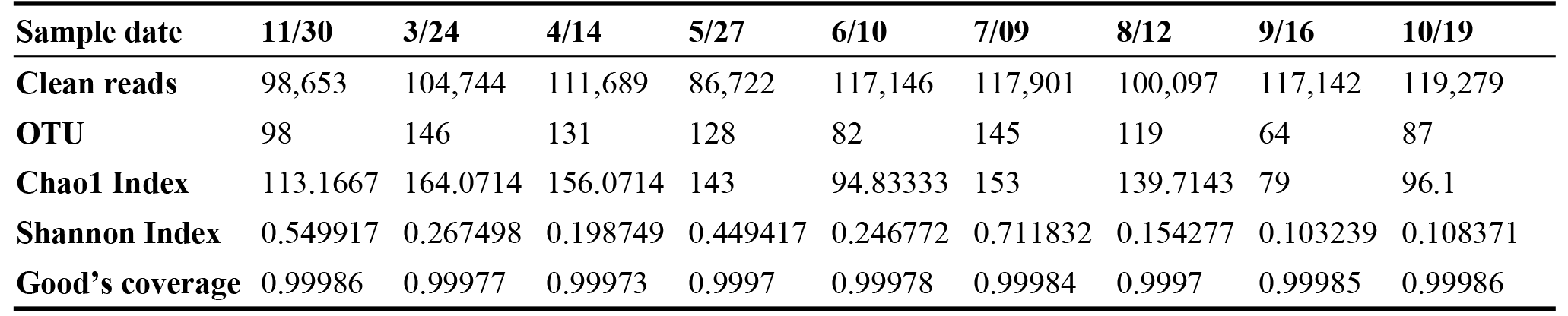
Distribution of OTUs among samples collected on different time.

| Sample date | 11/30 | 3/24 | 4/14 | 5/27 | 6/10 | 7/09 | 8/12 | 9/16 | 10/19 |
| --- | --- | --- | --- | --- | --- | --- | --- | --- | --- |
| Clean reads | 98,653 | 104,744 | 111,689 | 86,722 | 117,146 | 117,901 | 100,097 | 117,142 | 119,279 |
| OTU | 98 | 146 | 131 | 128 | 82 | 145 | 119 | 64 | 87 |
| Chao1 Index | 113.1667 | 164.0714 | 156.0714 | 143 | 94.83333 | 153 | 139.7143 | 79 | 96.1 |
| Shannon Index | 0.549917 | 0.267498 | 0.198749 | 0.449417 | 0.246772 | 0.711832 | 0.154277 | 0.103239 | 0.108371 |
| Good's coverage | 0.99986 | 0.99977 | 0.99973 | 0.9997 | 0.99978 | 0.99984 | 0.9997 | 0.99985 | 0.99986 |

### Major and minor OTUs

The 194 OTUs were divided into six groups based on their highest number of sequence reads at any time point (Table 2). Group 1 contains only one OTU, which is highly abundant with sequence reads over 30,000 in all samples at different time points. This OTU belongs to Buchnera, strongly suggesting that this is the principal endosymbiont of *Schlechtendalia chinensis*. Group 2, which comprises also highly abundant taxa, contains six OTUs with the highest numbers of reads between 100 to 2,000 per sample. The six OTUs belong to different taxonomic units including *Bacillus*, *Limnohabitans*, *Candidatus Cloacamonas*, *Comamonadaceae* and *Pseudomonas*. Group 3, defined here as medium-abundant OTUs, contains 38 OTUs with the highest numbers of reads between 10 to 100 in different samples. Twenty five OTUs had no blast hit, whereas the remaining OTUs corresponded to different taxa and are listed in the table 2. Group 4, which represent low abundant taxa, contains 47 OTUs with the highest read numbers between 5 and 10 in different samples. Twenty-eight OTUs had no blast hit, whereas the remaining OTUs corresponded to different taxonomic units. Group 5, comprisesinfrequent OTUs, contains 79 OTUs with the highest numbers of reads only between 2 to 5 in different samples, whereas group 6, referred as rare OTUs, contains 22 OTUs with the highest number of reads below 2.

**Table 2.** Groups of associated microbes with different levels of sequence reads. The lowest known level of taxonomy is given in the table. The numbers in parenthesis are the numbers of OTUs in that taxonomic unit.

| Group | Sequence reads | OTUs | OTU taxonomy |
| --- | --- | --- | --- |
| 1. Super abundant OTU | >30,000 | 1 | Buchnera (1) |
| 2. Highly abundant OTUs | <2,000 - 100 | 6 | Bacillus (1), Limnohabitans (1), Candidatus Cloacamonas (1), Comamonadaceae (1) Pseudomonas (2) |
| 3. Medium Abundant OTUs | <100 - 10 | 38 | Comamonas (1), Lactobacillales (1), Buchnera (2), Anaerolinaceae (1), Stenotrophomonas (1), Pirellulaceae (1), Rhodococcus (1), Streptococcus (1), Acinetobacter (1), Thermus (1), Crenarchaeota (1), Comamonadaceae (1), no blast hit (25) |
| 4. Low abundant OTUs | <10 - 5 | 47 | Xanthomonadaceae (1), Bacillus (2), Bacteria (1), Gemmataceae (1), Rhodobacter (1), Aminobacterium (1), Nitrosopumilus (1), Sphingomonas(1), Archaea (2), Pseudoxanthomonas (1), Buchnera (1), Anaerolinaceae (1), Methyloversatilis (1), Propionibacterium (1), Leuconostoc (1), Oxalobacteraceae (1), Anaerolinaceae (1), no blast hit (28) |
| 5. Infrequent OTUs | <5 - 2 | 79 | Buchnera (7), Alteromonadales (1), Comamonadaceae (7), Phormidium (1), Allochromatium (1), Rhizobium (1), Staphylococcus (1), Anoxybacillus kestanbolensis (1), Mycoplana (1), Archaea (1), Xenococcaceae (1), Enterobacteriaceae (3), Streptophyta (1), Bacillales (1), Pseudomonas stutzeri (1), Streptophyta (2), Acinetobacter (1), Halomonas (1), Methylobacteriaceae (1), Brevibacillus reuszeri (1), Roseomonas rosea(1), Caulobacteraceae (1), Burkholderiales (1), Rhizobiales (1), Wolbachia (1), no blast hit (39) |
| 6. Rare OUTs | <2 - 1 | 22 | Buchnera (5), Thauera (1), Brevundimonas diminuta (1), Enhydrobacter (1), no blast hit (14) |

### OTUs associated that differ between host plants

Samples collected in March, April, and November were isolated from the host moss, whereas the remaining samples were obtained from Sumac galls. There were 12 OTUs found exclusively in aphids collected from moss, and 11 OTUs were present exclusively in aphids collected from Sumac galls. The dynamic changes at different sampling time points of these OTUs specific to aphids feeding on the two different hosts are shown in Figs 1 and 2. The OTUs specific to aphids on different host plants were also listed in Table 3. There are five genera that were detected only in moss residing aphids. These, included *Wolbachia*, a genus from the family Gemmataceae, a genus from the family Pirellulaceae, *Phormidium*, and *Streptococcust*. And there are four genera (family) that were detected only in aphids feeding in Sumac galls, including *pGrfC26*, a genus from the family Xenococcaceae, a genus from the phylum Crenarchaeota, and *Nitrosopumilus*.

**Figure 1.**
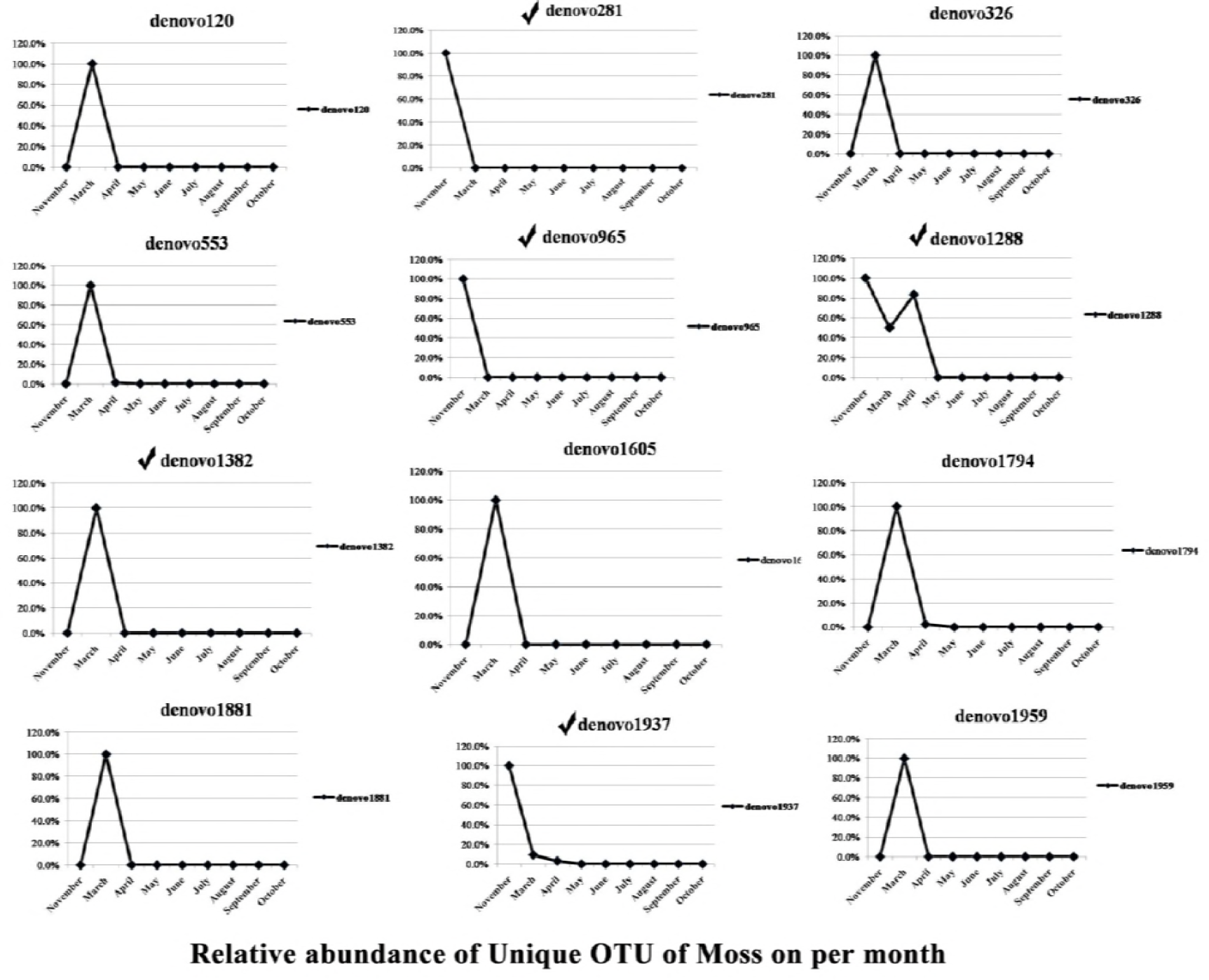
Dynamic changes at different sampling times of OTUs specific to aphids feeding on moss (from Nov to Apr). The symbol ✓ represents these OTUs that were classified into genus or family. The remaining OTUs had no blast hits.

**Figure 2.**
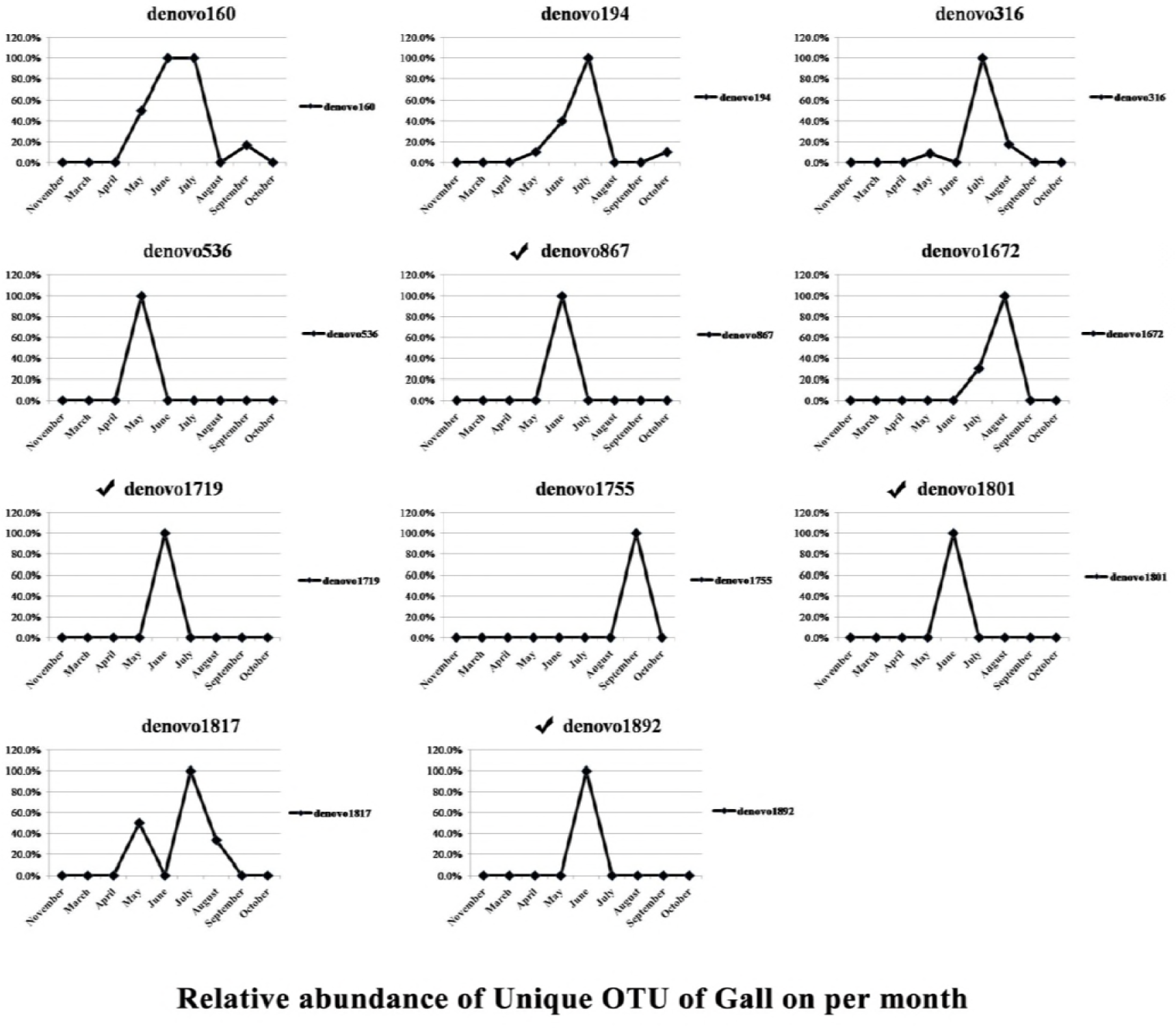
Dynamic changes at different sampling times of OTUs specific to aphids feeding in galls on Sumac trees (May to Oct when aphids lived in galls). The symbol ✓ indicates the OTUs that were classified into genus or family. The remaining OTUs had no blast hits.

**Table 3.** OTUs associated with aphids feeding on two different host plants.

| Host Plant | OTUs | Taxonomy (OTU name) |  |  |  |  |  |
| --- | --- | --- | --- | --- | --- | --- | --- |
| Exclusively in aphids from moss | 12 | No blast hit (denovo120) | *Wolbachia* (denovo1288) | Gemmataceae (denovo965) | No blast hit (denovo553) | Pirellulaceae (denovo281) | Comamonadaceae (denovo1605) |
|  |  | *Phormidium* (denovo1382) | No blast hit (denovo1794) | No blast hit (denovo1959) | No blast hit (denovo1881) | No blast hit (denovo326) | *Streptococcust* (denovo1937) |
| Exclusively in aphids from Sumac | 11 | *pGrfC26* (denovo1719) | Xenococcaceae (denovo1801) | *Buchnera* (denovo160) | No blast hit (denovo1817) | No blast hit (denovo1672) | No blast hit (denovo316) |
|  |  | No blast hit (denovo536) | *Crenarchaeota* (denovo867) | *Nitrosopumilus* (denovo1892) | *Streptophyta* (denovo194) | No blast hit (denovo1755) |  |
| Present in aphids from all samples | 23 | *Buchnera* (denovo528) | *Stenotrophomonas* (denovo2048) | *Buchnera* (denovo915) | *Acinetobacter* (denovo1435) | *Bacillus* (denovo187) | No blast hit (denovo780) |
|  |  | No blast hit (denovo2067) | *Pseudomonas stutzeri* (denovo739) | *Buchnera* (denovo2027) | *Pseudomonas* (denovo1118) | *Buchnera* (denovo1712) | *Buchnera* (denovo1556) |
|  |  | *Buchnera* (denovo753) | *Buchnera* (denovo496) | No blast hit (denovo1629) | *Streptophyta* (denovo743) | *Buchnera* (denovo155) | *Anoxybacillus kestanbolensis* (denovo1177) |
|  |  | Rhodococcus (denovo1058) | *Buchnera* (denovo104) | Comamonadaceae (denovo544) | *Buchnera* (denovo1489) | No blast hit (denovo1464) |  |

The OTUs were roughly equally present in aphids feeding on the two different hosts (Table 3). There are nine OTUs from different genera or families that were always present in aphids collected at different time points, including *Buchnera*, *Stenotrophomonas*, *Acinetobacter*, *Bacillus*, *Pseudomonas*, *Streptophyta*, *Anoxybacillus kestanbolensis*, Rhodococcus, and Comamonadaceae. The remaining OTUs were detected in aphids residing on either host plant, but their presence was not consistent at all time points.

### Beta diversity of microbiota during the whole life cycle of *S. chinensis*

A weighted UniFrac principal coordinates analysis (PCoA) was performed to compare the overall structure of microbiota from all samples based on the relative abundance of OTUs (Fig 3). There was an obvious separation between samples from different months even though some unevenly distributed presence. PC1, PC2 and PC3 accounted for 42.81%, 22.28% and 15.47% of total variations, respectively. This analysis showed differences in beta diversity resulted in separation between aphids feeding on moss and Sumac galls.

**Figure 3.**
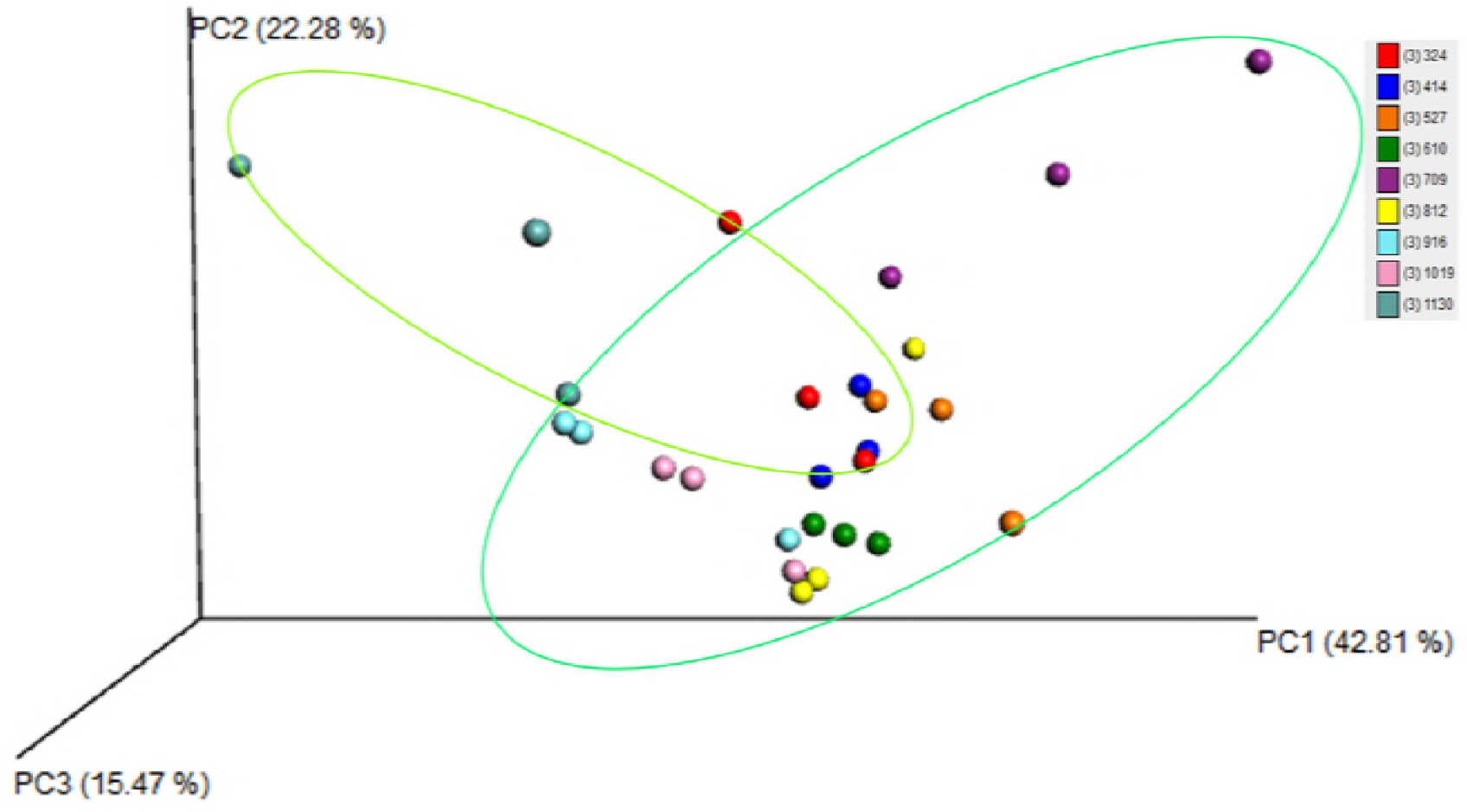
Weighted UniFrac principal coordinates analysis (PCoA) plots based on weighted UniFrac metric. Light green circle indicates data from aphids feeding on the moss. Dark green circle indicates data from aphids feeding in galls on Sumac trees.

## Discussion

The most abundant OTU was denovo496, which accounted for 80-99% of total sequence reads in all samples (S1 Table, S1 Fig). The taxonomy of denovo496 is *Buchnera*, which is an endosymbiont in all aphid species [3, 25–30]. (3, 25, 26, 27, 28, 29, 30) Since *Buchnera* is an obligate endosymbiont and is present in every aphid cell, it is not surprising that denovo496 was the most abundant microbe detected in our study. The abundance of sequence reads for *Buchnera* suggests that our sequence data were reliable and representative. Interestingly, there were 17 other OTUs corresponding to *Buchnera* in addition to denovo496 (S1 Table). The 17 OTUs were much less abundant with average sequence reads equal or below 16. However, many of these minor *Buchnera* OTUs, for example the OTU denovo915 (S1 Fig.), were consistently distributed in different samples, suggesting that these minor OTUs were not caused by sequence errors or other artifacts. The exact sources for these minor *Buchnera* OTUs remain to be determined. The minor OTUs could be due to minor alleles among *Buchnera* individuals in aphid cells.

The second-most abundant OTU was denovo743, which we detected in all samples across different time points. The percentages of denovo743 reads were below 0.8% of total reads in aphids feeding on moss, but the percentages reached as high as 7.5% of total reads in aphids living in galls. The taxonomy of denovo743 is *Streptophyta* from chloroplasts. Based on its taxonomy and abundance distribution, the sequences corresponding denovo743 were likely due to contamination of chloroplast DNA. It could be due to chloroplasts from gall tissues that stuck to the outside of aphid bodies. Alternatively, chloroplasts from gall tissues were sucked up to aphid guts.

Aside from *Buchnera*, other major bacteria associated with *S. chinensis* were from the genera *Bacillus*, *Limnohabitans*, *Candidatus Cloacamonas*, a genus from the family Comamonadacease, and *Pseudomonas* (Table 2, S1 Table). *Bacillus* is found commonly in aphids [31] and in other insect species [2, 32, 33, 34]. *Bacillus* protects the host from other bacterial and fungal colonization by producing antimicrobial compounds such as phenols [33, 35]. *Pseudomonas* bacteria colonize a wide range of ecological niches [27, 36], but it is not clear what kind of benefits this bacterium provides to aphids.

We detected 23 OTUs detected in all aphid samples collected from either moss or Sumac galls. Even though they were common to all samples, the abundance of these OTUs varied greatly from sample to sample exceptfor *Buchnera*. Therefore, it is not clear if these common OTUs are obligate or facultative and further research is needed to reveal their impact on the aphid host or aphid - plant interactions.

There were 12 OTUs unique to moss-residing aphid populations (Table 3). Typically, each OTU was detected at only one time point, either in November or March except the OTU denovo1288, which was found in Nov, Mar, and Apr samples (Fig 1). The presence of a specific symbiont at a specific time point indicated that these symbionts are likely facultative and play transient roles in the aphid life cycle. *Wolbachia* was among these moss-specific OTUs. *Wolbachia* has been previously found in aphids and it is a facultative symbionts that plays a role in the reproduction of its host insects [37, 38], as well as a role in host plant fitness [38]. Other symbionts specific to moss-residing aphids included bacteria from the families or genera Comamonadaceae, *Phormidium*, Pirellulaceae, Gemmataceae, and *Streptococcust*.

There were 11 OTUs specific to aphids associated with Sumac galls (Table 3). Most of these were present (or mainly present) in samples from a single time point (Fig 2), suggesting that they are likely facultative in their interaction with the host aphid. *Crenarchaeota* was among the symbionts unique in aphids collected from Sumac galls. *Crenarchaeota* symbionts have been reported to be involved nitrogen cycling [39]. *Nitrosopumilus* was another symbiont specific to aphids feeding in Sumac galls. *Nitrosopumilus* has been found to play roles in ammoxidation and CO2 assimilation in insects [40, 41].

In summary, a comprehensive survey was conducted on microbes associated with *S. chinensis* aphids on two different hosts at different time points. This survey identified a range of OTUs from highly abundant to rare microbes from different samples. We discovered both common and plant-specific symbionts. Identification of these symbionts provides a foundation for future studies to determine the roles of different symbionts on aphid physiology and insect - plant interactions.

## Acknowlegements

This study is funded by the Grant for Essential Scientific Research of National Non profit Institute (No. CAFYBB2014ZD005; CAFYBB2014QB019); and the National Natural Science Foundation of China (Grant No. U1402263; 31370651; 31372266; 31300019).

---

**Figure S1.**
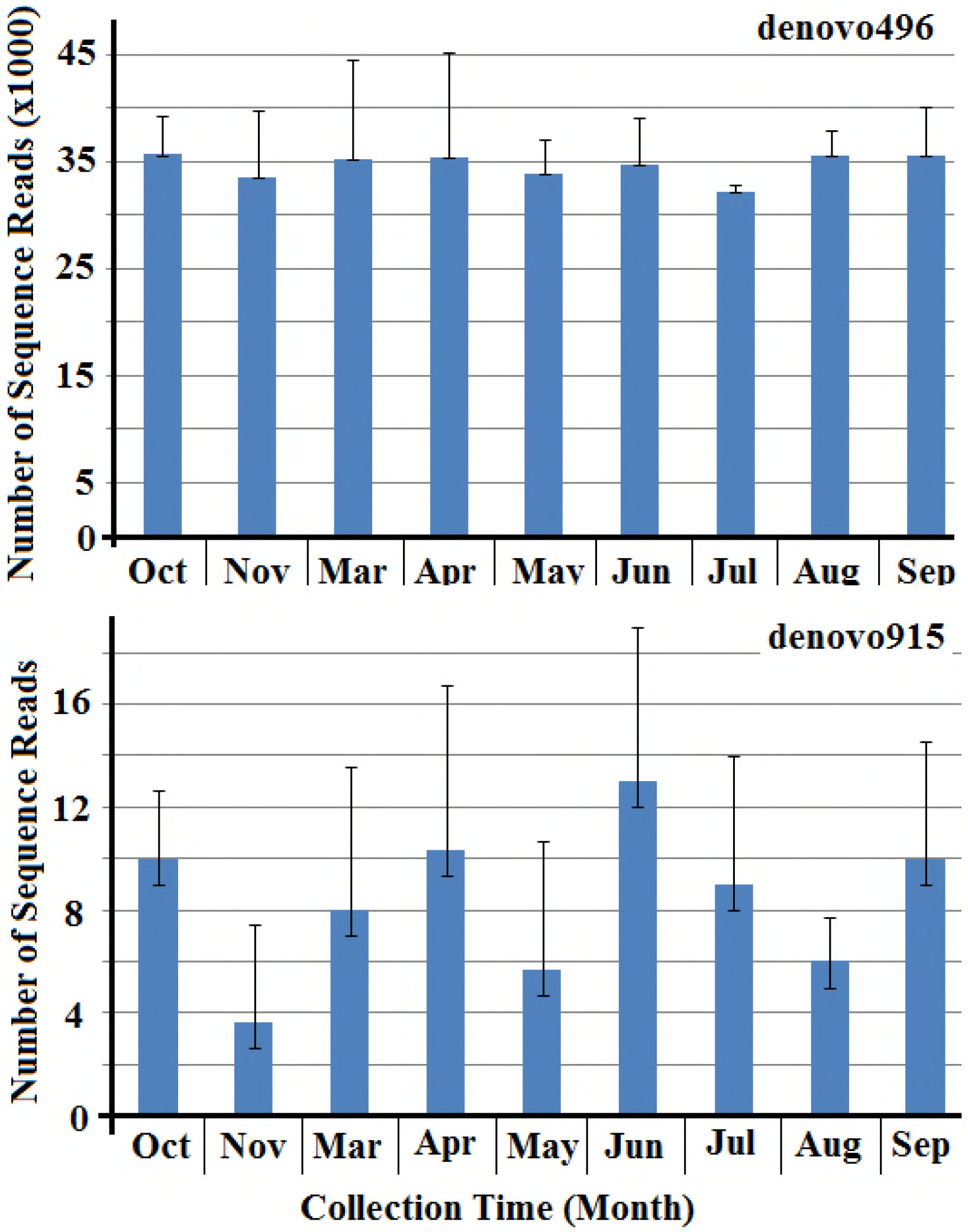
Abundance and variation of sequence reads for the two OTUs, denovo496 and denovo915, in different samples collected at different time points. The taxonomy of both OTUs correspond to *Buchnera*, an obligate endosymbiont. Standard errors are shown on the top of each bar.

## References

1. Dillon RJ, Dillon VM. The gut bacteria of insects: nonpathogenic interactions. Ann. Rev. Entomol. 2004; 49(1): 71–92. doi: 10.1146/annurev.ento.49.061802.123416

2. Bansal R, Hulbert S, Schemerhorn B, Reese JC, Whitworth RJ, Stuart JJ, et al. Hessian flyassociated bacteria: transmission, essentiality, and composition. PLoS ONE. 2011; 6:e23170. doi.: 10.1371/journal.pone.0023170

3. Shigenobu S, Wilson AC. Genomic revelations of a mutualism: the pea aphid and its obligate bacterial symbiont. Cell Mol. Life Sci. 2011; 68: 1297–1309. doi: 10.1007/s00018-011-0645-2

4. Gil R, Latorre A, Moya A. Bacterial endosymbionts of insects: insights from comparative genomics. Environ Microbiol. 2004; 6: 1109–1122. doi: 10.1111/j.1462-2920.2004.00691.x

5. Fellous S, Duron O, Rousset F. Adaptation dueto symbionts and conflicts between heritable agents of biological information. Nat Rev Gen. 2011; 12: 663. doi: 10.1038/nrg3028-c1

6. Douglas AE. Microbial Brokers of Insect-Plant Interactions Revisited. J Chem Ecol. 2013; 39(7): 952–961. doi: 10.1007/s10886-013-0308-x

7. Hansen AK, Moran NA. The impact of microbial symbionts on host plant utilization by herbivorous insects. Molecular Ecol. 2014; 23(6): 1473. doi: 10.1111/mec.12421

8. Oliver KM, Degnan PH, Burke GR, Moran NA. Facultative symbionts of aphids and the horizontal transfer of ecologically important traits. Annu Rev Entomol. 2010; 55: 247–266. doi: 10.1146/annurev-ento-112408-085305

9. Moran NA, Baumann P. Phylogenetics of cytoplasmically inherited microorganisms of arthropods. Trends Ecol. Evo. 1994; l9:15–20. doi:10.1016/0169-5347(94)90226-7

10. Moran NA, Russell JA, Koga R, Fukatsu T. Evolutionary Relationships of Three New Species of Enterobacteriaceae Living as Symbionts of Aphids and Other Insects. Appl Environ Microb. 2005; 71 (6): 3302–3310. doi: 10.1128/AEM.71.6.3302-3310.2005

11. Renoz F, Noël C, Errachid A, Foray V, Hance T. Infection dynamic of symbiotic bacteria in the pea aphid acyrthosiphon pisum gut and host immune response at the early steps in the infection process. PLoS ONE. 2015; 10(3): e0122099. doi: 10.1371/journal.pone.0122099

12. Leonardo T, Muiru GT. Facultative symbionts are associated with host plant specialization in pea aphid populations. Pro. Biol. Sci. 2003; 270(S2): S209–S212. doi: 10.1098/rsbl.2003.0064

13. Yang ZX, Chen XM, Feng Y, Chen H. Morphology of the antennal sensilla of Rhus gall aphids (Hemiptera: Aphidoidea: Pemphiginae): A comparative analysis of five genera. Zootaxa. 2009; 2204: 48–54.

14. Yang ZX, Chen XM, Havill NP, Feng Y, Chen H. Phylogeny of Rhus gall aphids (Hemiptera: Pemphigidae) based on combined molecular analysis of nuclear EF1a and mitochondrial COII genes. Entomological Science. 2010; 13: 351–357. doi: 10.1111/j.1479-8298.2010.00391.x

15. Stroyan HG. Aphids. In McGraw-Hill Encyclopedia of Science and Technology, 8th Edition, ISBN 0-07-911504-7; 1997.

16. Shao SX, Yang ZX, Chen XM. Gall Development and Clone Dynamics of the Galling Aphid Schlechtendalia chinensis (Hemiptera: Pemphigidae). J. Econ. Entomol. 2013; 106(4):1628–1637. doi: 10.1603/EC13114

17. Liu P, Yang ZX, Chen XM, Foottit RG. The Effect of the Gall-Forming Aphid Schlechtendalia chinensis (Hemiptera: Aphididae) on Leaf Wing Ontogenesis in Rhus chinensis (Sapindales: Anacardiaceae). Ann. Entomol. Soc. Am. 2014; 107(1):242–250. doi: 0.1603/AN13118

18. Ekblom R, Galindo J. Applications of next generation sequencing in molecular ecology of non model organisms. Heredity. 2011; 107:1–15. doi: 10.1038/hdy.2010.152

19. Magoč T, Salzberg SL. FLASH: fast length adjustment of short reads to improve genome assemblies. Bioinformatics. 2011; 27.21: 2957–2963. doi: 10.1093/bioinformatics/btr50720.

20. Caporaso JG, Kuczynski J, Stombaugh J, Bittinger K, Bushman FD, Costello EK et al. QIIME allows analysis of high-throughput community sequencing data. Nature methods. 2010; 7.5: 335–336. doi: 10.1038/nmeth.f.303

21. Edgar RC. UPARSE: highly accurate OTU sequences from microbial amplicon reads. Nature methods. 2013; 10.10: 996–998. doi: 10.1038/nmeth.2604

22. Wang Q, Garrity GM, Tiedje JM., Cole JR. Naive Bayesian classifier for rapid assignment of rRNA sequences into the new bacterial taxonomy. Appl. Environ. Microb. 2007; 73: 5261–5267. doi: 10.1128/AEM.00062-07

23. DeSantis TZ, Hugenholtz P, Larsen N, Rojas M, Brodie EL, Keller K, et al. Green genes, a chimera-checked 16S rRNA gene database and workbench compatible with ARB. Appl. Environ. Microb. 2006; 72.7: 5069–5072. doi: 10.1128/AEM.03006-05

24. Huson D, Mitra S, Ruscheweyh HJ, Weber N, Schuster SC. Integrative analysis of environmental sequences using M EGAN4. Genome Res. 2011; 21: 1552–1560. doi: 10.1101/gr.120618.111

25. Baumann P, Baumann L, Lai CY, Rouhbakhsh D, Moran NA, Clark MA. Genetics, physiology, and evolutionary relationships of the genus Buchnera: Intracellular symbionts of aphids. Annu. Rev. Microbiol. 1995; 49: 55–94. doi: 10.1146/annurev.mi.49.100195.000415

26. Baumann P, Lai CY, Baumann L, Rouhbakhsh D, Moran NA, Clark MA. Mutualisticassociations of aphids and prokaryotes; biology of the genus Buchnera. Appl Env Microbiol. 1995; 61:1–7.

27. Bansal R, Mian MA, Michel AP. Microbiome diversity of Aphis glycines with extensive superinfection in native and invasive populations. Environ Microbiol Rep. 2014; 6(1):57–69. doi: 10.1111/1758-2229.12108

28. Liu S, Chougule NP, Vijayendran D, Bonning BC. Deep sequencing of the transcriptomes of soybean aphid and associated endosymbionts. PloS ONE. 2012; 7:e45161. doi: 10.1371/journal.pone.0045161

29. Russell CW, Bouvaine S, Newell P D, Douglas AE. Shared metabolic pathways in a coevolved insect-bacterial symbiosis. Appl. Environ. Microb. 2013; 79(19): 6117–6123. doi: 10.1128/AEM.01543-13

30. Shigenobu S, Stern DL. Aphids evolved novel secreted proteins for symbiosis with bacterial endosymbiont. P Roy Soc B-Biol Sci. 2013; 280(1750): 20121952.

31. Ateyyat M. Culturable bacteria associated with the guts of pea aphid, Acyrthosiphum pisum (Homoptera: Aphididae). J Entomol. 2008; 5: 167–175. doi: 10.3923/je.2008.167.175

32. Behar A, Jurkevitch E, Yuval B. Bringing back the fruit into fruit fly-bacteria interactions. Mol Ecol. 2008; 17: 1375–1386. doi: 10.1111/j.1365-294X.2008.03674.x

33. Blackburn MB, Gunderson-Rindal DE, Weber DC, Martin PAW, Farrar RRJ. Enteric bacteria of field-collected Colorado potato beetle larvae inhibit growth of the entomopathogens Photorhabdus temperata and Beauveria bassiana. Biol Control. 2008; 46: 434–441. doi: 10.1016/j.biocontrol.2008.05.005

34. Feng W, Wang XQ, Zhou W, Liu GY, Wan YJ. Isolation and characterization of lipase-producing bacteria in the intestine of the silkworm, Bombyx mori, reared on different forage. J Insect Sci. 2011; 11: 1–10. doi: 10.1673/031.011.13501

35. Dillon RJ, Vennard CT, Buckling A, Charnley AK. Diversity of locust gut bacteria protects agains pathogen invasion. Ecol Lett. 2005; 8: 1291–1298. doi: 10.1111/j.1461-0248.2005.00828.x

36. Chevalier S, Bouffartigues E, Bodilis J, Maillot O, Lesouhaitier O, Feuilloley MG, Cornelis P. Structure, function and regulation of Pseudomonas aeruginosa porins. FEMS Microbiol Rev. 2017; fux020. hdoi: 10.1093/femsre/fux020

37. Augustinos AA, Santos-Garcia D, Dionyssopoulou E, Moreira M, Papapanagiotou A, Scarvelakis M. et al. Detection and characterization of Wolbachia infections in natural populations of aphids: is the hidden diversity fully unraveled? PLoS ONE. 2011; 6: e28695. doi: 10.1371/journal.pone.0028695

38. Zytynska SE, Weisser WW. The natural occurrence of secondary bacterial symbionts in aphids. Ecol entomol. 2016; 41(1):13–26. doi: 10.1111/een.12281

39. Nicol GW, Schleper C. Ammonia-oxidising Crenarchaeota: important players in the nitrogen cycle? Trends Microbiol. 2006; 14(5): 207–212. doi: 10.1016/j.tim.2006.03.004

40. Jia ZJ, Conrad R. Bacteria rather than Archaea dominate microbial ammonia oxidation in an agricultural soil. Environ Microbiol. 2009; 11 (7): 1658–1671. doi: 10.1111/j.1462-2920.2009.01891.x

41. Xia WW, Zhang CX, Zeng XW, Feng YZ, Weng JH, Lin XG. et al. 2011. Autotrophic growth of nitrifying community in an agricultural soil. The ISME Journal 5 (7):1226–1236. https://doi.org/10.1038/ismej.2011.5

